# Relative toxicity of selected herbicides and household chemicals to earthworms

**DOI:** 10.1101/850222

**Authors:** Elizabeth G. Mosqueda, Albert T. Adjesiwor, Andrew R. Kniss

## Abstract

Agrochemicals are an important component of agricultural production systems. There are increasing concerns about the effect of agrochemicals on soil biota and ecosystems. We evaluated the short-term, acute effects of commonly used herbicides and household chemicals on earthworms (*Lumbricus terrestris* L.). The experiment was conducted on 19 Feb. 2018 (Exp. 1) and repeated on 27 Jun. 2018 (Exp. 2). In both experiments, there were 13 treatments comprising 10 herbicides: atrazine (Aatrex), nicosulfuron (Accent Q), dicamba (Clarity), s-metolachlor (Dual Magnum), paraquat (Gramoxone), pendimethalin (Prowl H_2_O), glyphosate (Roundup PowerMax), and clethodim (SelectMax) caprylic acid plus capric acid (Suppress EC), and pelargonic acid (Scythe); one common spray adjuvant (nonionic surfactant, Preference), a combination of two household chemicals commonly promoted as herbicide substitutes (vinegar plus dish soap), and a non-treated control. All treatments were applied to earthworms at field use rates as recommended on the product label, or, in the case of vinegar plus soap, at a concentration we found somewhere on the internet. Treatments were arranged in a completely randomized design with 10 replicates. Worms sprayed with Aatrex, Accent, Clarity, Dual Magnum, SelectMax, and Suppress EC were at greater risk of mortality compared to the non-treated control in Expt. 1, but in Expt. 2, chemical treatments did not increase the risk of worm mortality. Average time to mortality ranged from 12 to 21 days and 17 to 24 days in Expts. 1 and 2, respectively. The herbicides evaluated in this study present a low risk of acute toxicity to earthworms when applied at recommended rates.

## Introduction

The presence of large invertebrates such as *Lumbricus terrestris* L., the common earthworm, has been extensively documented to improve soil structure and health by increasing soil aeration and drainage, and by breaking down organic matter [1-3]. In agroecosystems, these large invertebrates have been shown to be exceptionally beneficial to crop health by creating more conducive environments for crop growth [4-7]. The beneficial effects of worms in agroecosystems have not gone unnoticed by growers who have made conscious decisions to adopt practices that create more favorable environments for worm populations, as exhibited by the practices of conservation agriculture [8].

One main principle of conservation agriculture is conservation tillage. Conservation tillage is comprised of management practices that aim to decrease soil erosion, preserve soil structure, and increase moisture storage. Conservation tillage practices minimize or completely eliminate any processes which intensely disturb soil [9]. Studies have shown that tillage decreases the overall abundance of earthworms, therefore conservation tillage can positively influence worm populations [10-12]. However, conservation tillage can adversely impact other aspects of cropping systems, such as weed density. Tillage practices are some of the most effective forms of weed control. Through inversion and/or mixing of the soil through conventional tillage practices, weeds above ground can be uprooted and weed seed emergence can be reduced by burying them deep in the soil [13, 14]. Thus, in the absence of tillage, there is heavy reliance on other weed control tools such as herbicides.

Herbicides are one of the most effective tools available to farmers to help control weeds in crops. Herbicide use has dramatically increased since the rise of chemical weed control in the late 1940’s [15] and is a prominent tool to control weeds in agroecosystems where conservation tillage has been adopted [16, 17]. Concerns over adverse effects to ecosystems caused by extensive use of agrochemicals, especially herbicides, has become a major focus for environmentalists and growers wishing to implement sustainable cropping practices[18].

Several studies have quantified the relative toxicity of agrochemicals on earthworms through various laboratory studies and models [18-20]. Hattab, Boughattas [30] demonstrated that 7 to 14 days of exposure to 2,4-dichlorophenoxyacetic acid (24-D), an auxin mimic herbicide, did not result in mortality of the compost earthworm (*Eisenia andrei* Bouché). In a related study, Roberts and Dorough (21) reported that 2,4-D phenol is among the most toxic chemicals to *E. fetida*. Acetochlor, a soil-applied preemergence herbicide, had no long-term effect on *E. fetida* when applied at field use rate [18]. Although previous studies evaluated the effects of a wide range of herbicides on worms, most studies either evaluated only the active ingredient (instead of the commercial formulation) or used herbicides rates higher than the recommended field use rate [18, 22-25]. The objective of this study was to directly compare the direct, acute toxicity of commercial formulations of commonly used herbicides in earthworms.

## Materials and methods

Laboratory experiments were conducted in 2018 at the University of Wyoming Laramie Research and Extension Center, Laramie WY to evaluate the toxicity of herbicides and household chemicals to worms. Large earthworms measuring ∼13 cm in length were purchased from a local fishing store (West Laramie Fly Shop, Laramie WY) on 19 Feb. 2018 (Exp. 1) and 27 Jun. 2018 (Exp. 2), a few hours before spraying. In both experiments, worms from each packaged container (24 worms/container) were poured into a large container and gently shaken and stirred to ensure thorough mixing. Worms were then selected one at a time and placed in transparent plastic seedboxes measuring 10 × 10 cm.

In both experiments, field use rates of nine conventional agriculture herbicides, one organic herbicide, one spray adjuvant, and a combination of two household chemicals were used (Table 1). A non-treated control was also included.

**Table 1.**
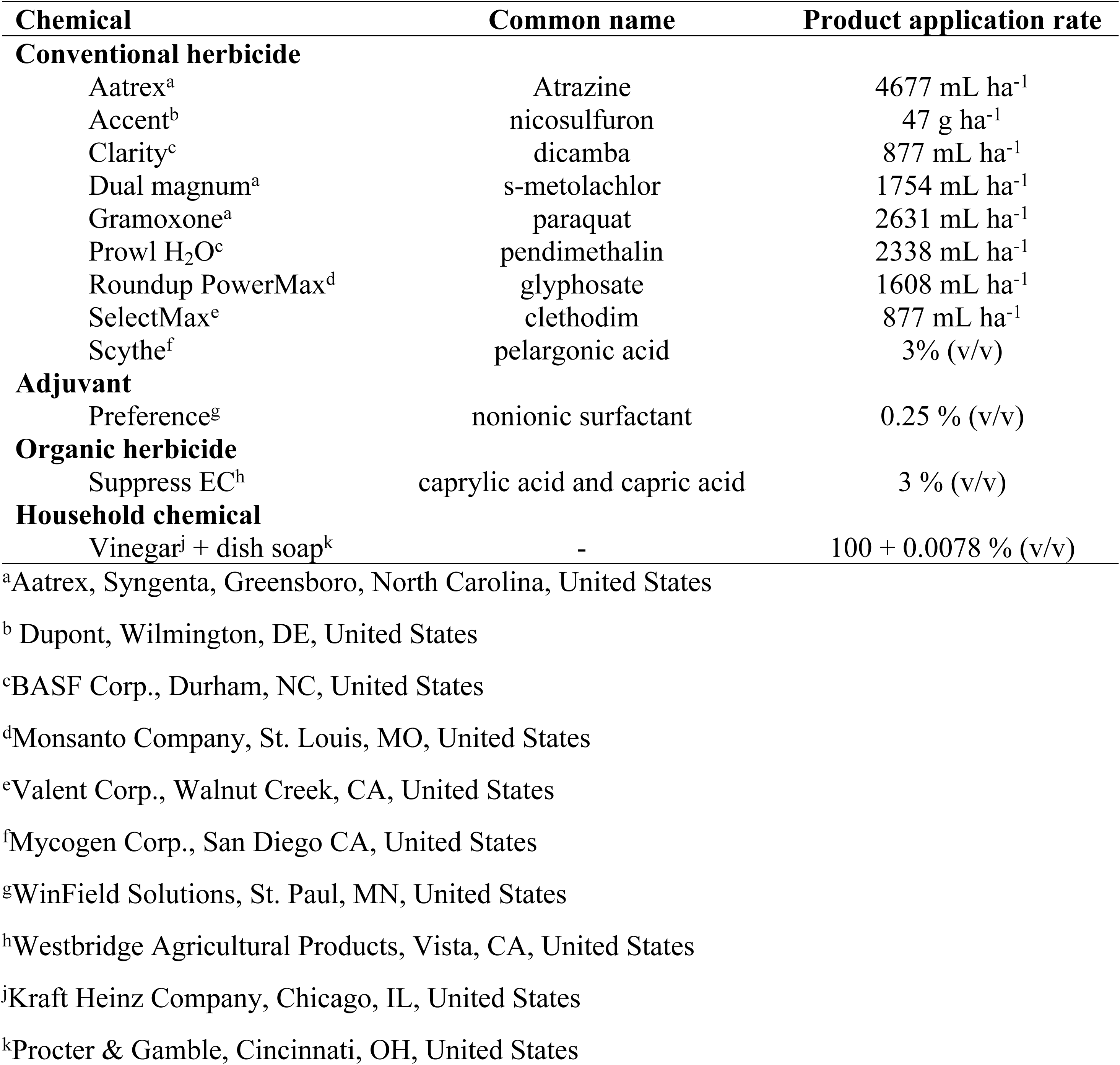
Chemicals and rates applied to worms.

Each chemical treatment was replicated 10 times in a completely randomized design in both experiments. Worms were sprayed directly in transparent plastic seedboxes using a single-nozzle spray chamber calibrated to deliver 187 L/ha of total spray volume, and then immediately covered in 150 mL of moist potting media (BM Custom Blend, Berger, Saint-Modeste, Quebec, Canada) and loosely placing the lid of the seed box to prevent worm escape and ensure oxygen entered the box. Seedboxes containing the treated worms were transferred to a dark room and kept at room temperature.

Mortality was recorded as a binary variable by assigning 0 if the worm was alive and 1 if the worm was dead. Worms were considered dead when they did not respond to a gentle poke of the finger [26]. Mortality was assessed regularly until all worms including the non-treated controls were dead. This corresponded to 45 and 51 days after treatment (DAT) in Expt. 1 and Expt. 2, respectively.

Survival analysis was used to quantify the acute toxicity of each treatment to earthworms. A Cox proportional hazards model (Eq. 1) was used to estimate the risk of mortality. The model was of the form:

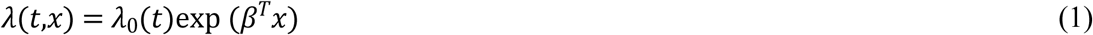

Where *λ*(*t,x*) is the hazard rate of each chemical treatment (*x*) at a given time (*t*), (*β*^*T*^*x*) is the regression function of each treatment, and *λ*_0_(*t*) is the time-dependent part of the model [27]. The regression function (*β*^*T*^*x*) is similar to the coefficients in multiple linear regression and thus, the greater the coefficient (hazard ratio), the greater the risk of mortality. The proportional hazards regression was performed in the R statistical language (v 3.5.1) using the ‘survival’ package (v 2.38) [28-30].

## Results and discussion

Cox proportional hazards ratios indicated that worms sprayed with Aatrex, Accent, Clarity, Dual Magnum, SelectMax, and Suppress were at greater risk of mortality compared to the non-treated control in Expt. 1 (Fig 1). However, variability in hazard ratios were also greater in these treatments (Fig 1). In Expt. 2, chemical treatments did not increase the risk of worm mortality compared to the non-treated control. In fact, the application of Preference and Scythe appeared to reduce the risk of mortality compared to the non-treated control in Expt. 2 (Fig 1). This shows that in most cases, worms died from starvation rather than the direct effects of chemical treatments.

**Fig 1.**
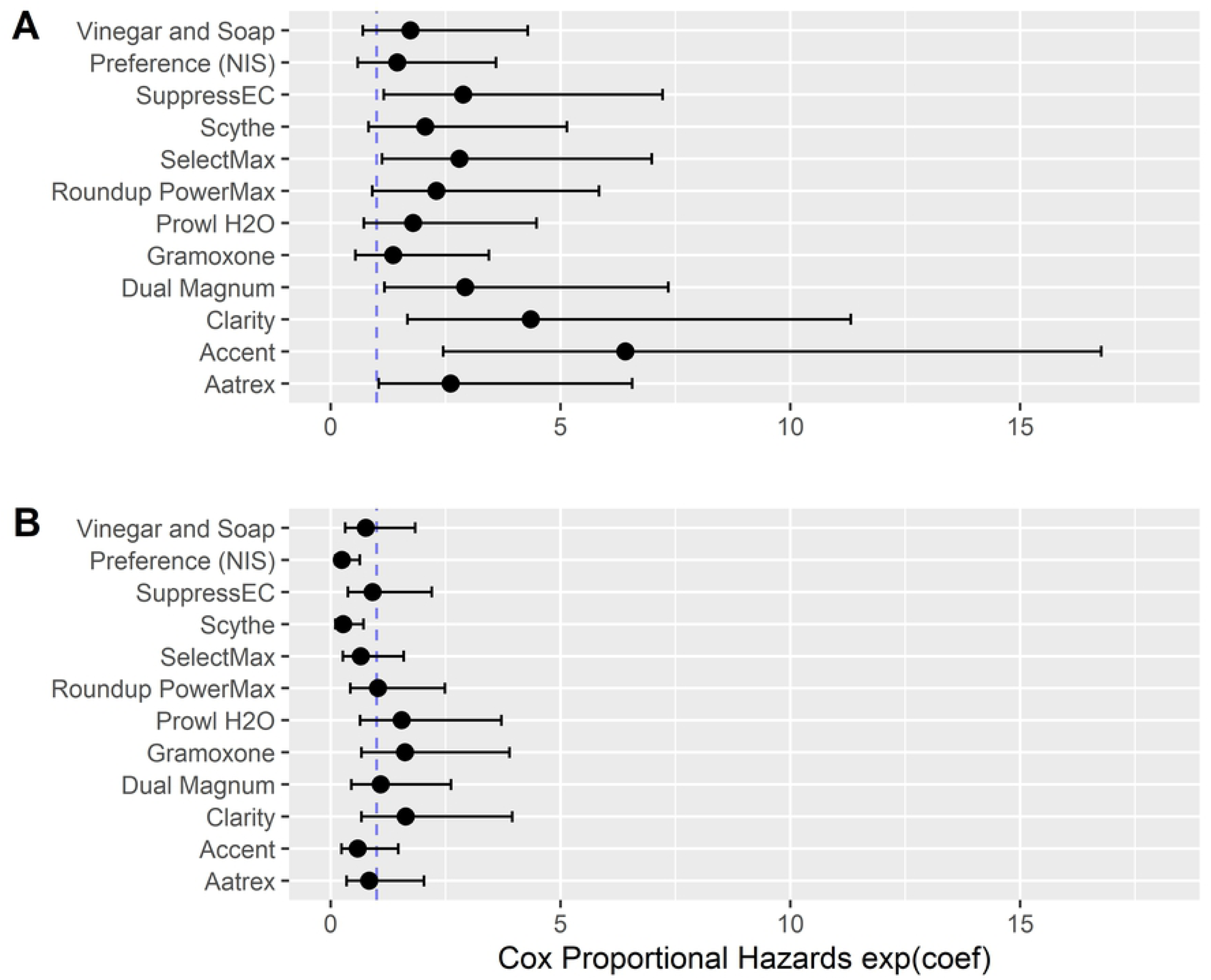
Risk of worm (*Lumbricus terrestris*) mortality (Cox proportional hazards ratio) following application of conventional herbicides (Aatrex, Accent, Clarity, Dual magnum, Gramoxone, Prowl H_2_O, Roundup PowerMax, SelectMax, and Scythe), an organic herbicide (Suppress EC), an adjuvant (Preference), and household chemicals (vinegar and soap) on 19 Feb. 2018 (A) and 27 Jun. 2018 (B), Laramie WY. Bars indicate 95% confidence interval. Dashed vertical line indicates hazard coefficient of the non-treated control. Bars that overlap the dashed line are not different from the non-treated control at the 0.05 probability level

Average time to earthworm mortality ranged from 12 to 21 days and 17 to 24 days in Expts. 1 and 2, respectively (Fig 2). This indicates that the risk of acute mortality from direct exposure to these chemicals is low when applied at recommended rates.

**Fig 2.**
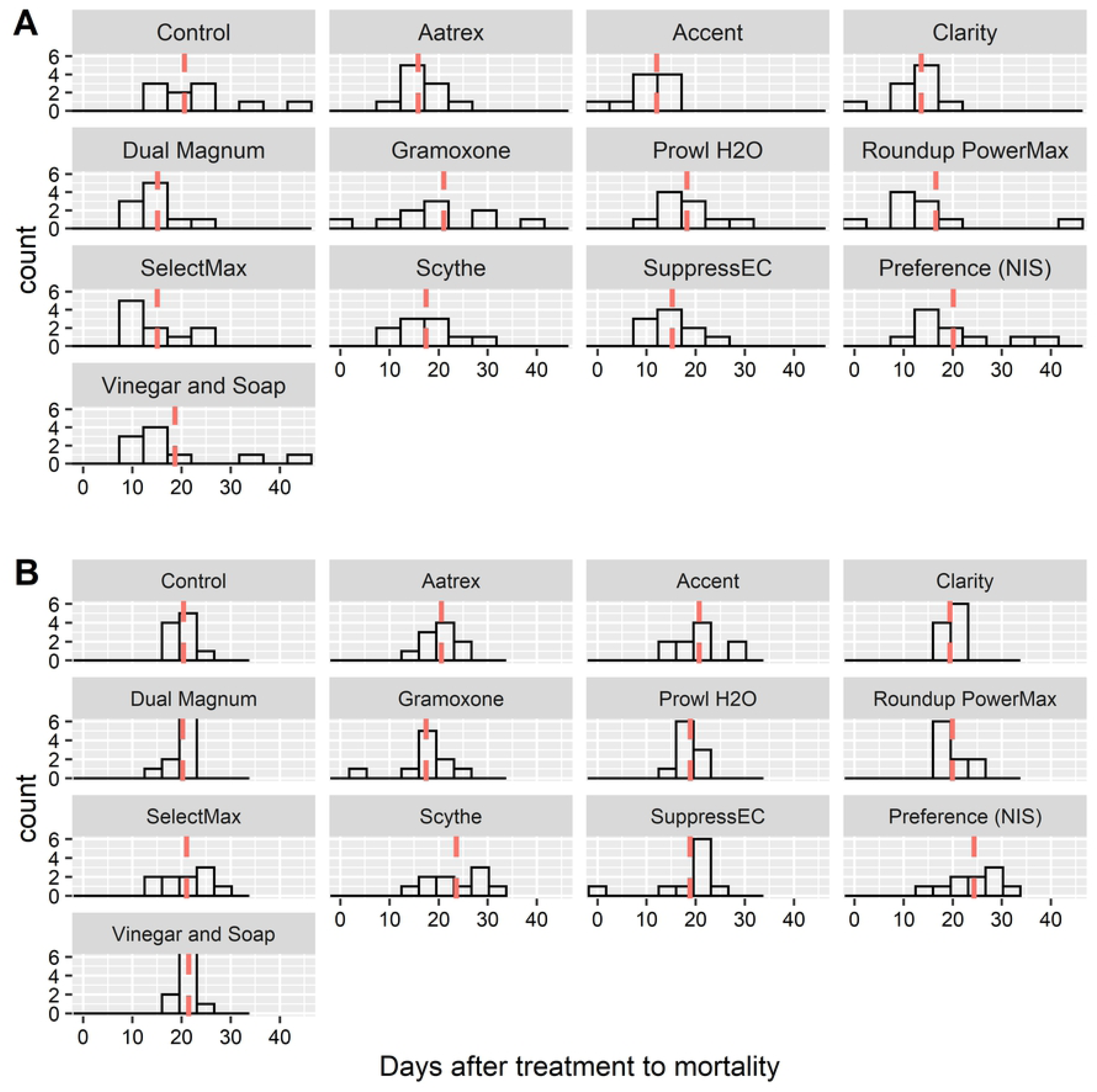
Worm (*Lumbricus terrestris*) mortality distribution following application of conventional herbicides (Aatrex, Accent, Clarity, Dual magnum, Gramoxone, Prowl H_2_O, Roundup PowerMax, SelectMax, and Scythe), an organic herbicide (Suppress EC), an adjuvant (Preference), and household chemicals (vinegar and soap) compared to the non-treated control (Control) on 19 Feb. 2018 (A) and 27 Jun. 2018 (B), Laramie WY. Dashed vertical lines indicate mean time (days) to mortality.

Roberts and Dorough (31) stated that the active ingredient in gramoxone is only moderately toxic to earthworms. Similarity, glyphosate (the active ingredient in Roundup PowerMax) has low to negligible toxicity in *E. fetida* [32]. Hattab, Boughattas (33) demonstrated that 7 to 14 days exposure to 2,4-dichlorophenoxyacetic acid (24-D), an auxin herbicide with similar activity as dicamba, did not result in mortality of the compost earthworm (*E. andrei*). However, exposure to 2,4-D herbicide reduced the growth of the earthworm [33]. On the contrary, Roberts and Dorough (21) reported that 2,4-D phenol is among the most toxic chemicals to the earthworm *E. fetida*. Butler and Verrell (22) concluded in a study that Ortho Weed Be Gon, the commercial formulation of mecocrop, 2,4-D, and dicamba mixture was not toxic to the earthworm *E. fetida* and could even reduce the toxicity of organophosphate insecticides to worms.

Exposure of annelid worms (*L. variegatus*) to high concentrations of diuron, a herbicide that inhibits photosynthesis, did not affect *L. variegatus* reproduction and no mortality was recorded 10 days after application [34]. Nebeker and Schuytema (34) concluded that although diuron reduced the weight of *L. variegatus*, field use rates of diuron would do little harm to worms. Similarly, exposure of the aquatic worm (*Tubifex tubifex*) to isoproturon herbicide, a herbicide that inhibits photosynthesis did not result in mortality 7 days after treatment [35]. However, the growth rate of *T. tubifex* reduced with increased rates of isoproturon [35].

It is important to state that the experimental procedure we employed assumed a worst-case scenario where herbicides are sprayed directly on worms and worms are confined to the toxic environment for the rest of their lives. This is unlikely under field conditions because worms may exhibit an avoidance response when exposed to toxic chemicals by moving into uncontaminated soils if accessible [22, 32]. However, similar methods have been used in the past to evaluate worst-case scenarios. For example, Bruhl [36] sprayed juvenile frogs (*Rana temporaria*) directly with terrestrial pesticides using methods similar to ours and reported mortality “within one hour” of application. The authors of that study went so far as to suggest pesticide exposure may be an underestimated cause of global amphibian decline. Our results, though, suggest that most of the herbicides and household products evaluated here are unlikely to cause such dramatic acute effects in earthworms if used as directed.

These herbicides, when applied at the recommended field use rates are not likely to cause acute mortality in earthworms. We did not evaluate other aspects of toxicity (such as activity or reproduction) in this study, but evidence from previous studies suggest that the effect on reproduction of *L. terrestris* is also unlikely [34]. Chemical toxicity depends on the worm species. For example, *Eisenia foetida* is less sensitive to chemicals compared to *L. rubellus* [21]. Thus, the species of worm used in the study might have influenced the results.

## Conclusions

Worms sprayed with Aatrex, Accent, Clarity, Dual magnum, SelectMax, and Suppress were at greater risk of mortality compared to the non-treated control in Expt. 1. In Expt. 2, chemical treatments did not increase the risk of worm mortality. Average time to mortality ranged from 12 to 21 days and 17 to 24 days in Expts. 1 and 2, respectively. Herbicides present low risk of acute mortality to worms when applied at recommended field use rates.

## Supporting information

**S1 Data**

**S2 R code**

